# Structural basis of the human Scribble-Vangl2 association in health and disease

**DOI:** 10.1101/2020.10.07.330712

**Authors:** Jing Yuan How, Rebecca K. Stephens, Krystle Y.B. Lim, Patrick O. Humbert, Marc Kvansakul

## Abstract

Scribble is a critical cell polarity regulator that has been shown to work as either an oncogene or tumor suppressor in a context dependent manner, and also impacts cell migration, tissue architecture and immunity. Mutations in Scribble lead to neural tube defects in mice and humans, which has been attributed to a loss of interaction with the planar cell polarity regulator Vangl2. We show that the Scribble PDZ domains 1, 2 and 3 are able to interact with the C-terminal PDZ binding motif of Vangl2 and have now determined crystal structures of these Scribble PDZ domains bound to the Vangl2 peptide. Mapping of mammalian neural tube defect mutations reveal that mutations located distal to the canonical PDZ domain ligand binding groove can not only ablate binding to Vangl2 but also disrupt binding to multiple other signaling regulators. Our findings suggest that PDZ-associated neural tube defect mutations in Scribble may not simply act in a Vangl2 dependent manner but as broad-spectrum loss of function mutants by disrupting the global Scribble-mediated interaction network.

## Introduction

The establishment of cell polarity, defined as the asymmetric distribution of proteins, carbohydrates and lipids within the cell, is a crucial process for the organization and development of all animal tissues (Nelson, 2003) In multicellular organisms, four major different types of polarity can be distinguished: apico-basal cell polarity, asymmetric cell division, front-rear cell polarity and planar cell polarity (Dow and Humbert, 2007).

Planar cell polarity is a critical feature of the extension and axial elongation of tissues, establishing uniformly directional information through coordinating polarity across cells within the tissue plane (Butler and Wallingford, 2017). In mammals, planar cell polarity is essential during embryonic developmental stages, where its dysregulation during this time results in the neural tube failing to close and subsequent congenital malformations, including spina bifida and craniorachischisis, a severe neural tube defect in which the midbrain, hindbrain and the entire spinal region remain open (Nikolopoulou et al., 2017). Planar cell polarity deregulation is further implicated in defects in wound closure, a process that requires coordination of multiple cells across a tissue to migrate and proliferate together, as well as the early stages of cancer (Munoz-Soriano et al., 2012).

The Scribble protein is a highly conserved cell polarity regulator comprising 16 Leucine Rich Repeats and four PSD-95/Discs-large/ZO-1 (PDZ) domains and belongs to the LAP family of proteins (Bonello and Peifer, 2019, Santoni et al., 2020). Together with Discs Large (Dlg) and Lethal Giant Larvae (Lgl), Scribble acts to establish and maintain tissue homeostasis, as well as directed cell migration and tissue growth (Stephens et al., 2018). Whilst Scribble was originally characterized as a cell polarity regulator and tumour suppressor in the vinegar fly *Drosophila melanogaster* (Bilder et al., 2000), its role in planar cell polarity was first identified in the mouse, where mutation to Scribble result in a severe, embryonic lethal form of neural tube defect known as craniorachischisis (Murdoch et al., 2003)(Reviewed in Milgrom-Hoffman and Humbert, 2018). Importantly, Scribble mutations leading to neural tube defects have also been identified in humans, further highlighting its importance as a candidate gene to study planar cell polarity regulation and developmental defects (Robinson et al., 2012, Lei et al., 2013).

Scribble modulates numerous cellular processes through its many protein-protein interactions, which are largely mediated by Scribble’s four PDZ domains (Stephens et al., 2018). These PDZ domains bind C-terminally located PDZ-binding motifs (PBMs) on specific interactors in a selective manner, whilst individual Scribble PDZ domains display overlapping specificities for particular ligands (Stephens et al., 2018).

One such Scribble-interactor is Vangl2 (mammalian homologue of *Drosophila* Van Gogh), a transmembrane protein identified as a critical component of the planar cell polarity pathway (Bailly et al., 2018). Similarly to Scribble, Vangl2 is essential in early embryonic development where its loss leads to neural tube defects as well as disruption in the cochleae inner ear hair bundle orientation (Murdoch et al., 2001, Montcouquiol et al., 2003). A number of mutations in human Vangl2 have also been shown to be associated to neural tube closure defects (Kibar et al., 2011) with recent studies implicating disruption of planar cell polarity as the causative factor for these mutations (Humphries et al., 2020). Importantly, Vangl2 genetically interacts with Scribble to regulate planar cell polarity processes in the mouse (Montcouquiol et al., 2003, Murdoch et al., 2003, Phillips et al., 2007, Yates et al., 2013). Furthermore, Vangl2 has been shown to bind to Scribble PDZ domains both *in vivo* and *in vitro* (Montcouquiol et al., 2003, Kallay et al., 2006, Montcouquiol et al., 2006) and its loss has been shown to lead to Vangl2 mislocalisation in mouse and zebrafish models (Phillips et al., 2007, Navajas Acedo et al., 2019). Specifically, Vangl2 was shown to engage Scribble PDZ2 and 3 domains via a C-terminal PBM (Kallay et al., 2006, Montcouquiol et al., 2006).

However, a more detailed molecular understanding of the relationship between Scribble and Vangl2 has been missing. We now report a systematic evaluation of Vangl2 PBM interactions with Scribble PDZ domains, together with structural analysis of all identified Scribble-Vangl2 interactions. Furthermore, using mutagenesis we biochemically interrogate the consequences of reported Scribble mutations associated with neural tube defects, thus providing a mechanistic basis to understand the interplay between Scribble and Vangl2 during correct establishment of planar cell polarity as well as in a disease state such as during neural tube closure defects.

## Results

### Scribble PDZ domains show interactions with Vangl2

To understand the molecular basis for Scribble:Vangl2 interactions and its consequences on correct tissue development, in particular neural tube closure, we systematically examined interactions between individual Scribble PDZ domains with the C-terminal PBM of Vangl2 (RLQSETSV) using isothermal titration calorimetry. We unexpectedly found that Scribble PDZ1, 2 and 3 domains are able to all interact with the PBM of Vangl2, with affinities ranging from 24-45 μM (Figure 1), whereas Scribble PDZ4 did not show any detectable affinity. As expected, a mutant Vangl2 peptide (RLQSEASA) did not show any binding to Scribble PDZ domains (Table 1). These findings are in contrast to previous data indicating that Scribble engages Vangl2 only via its PDZ2 and 3 domains (Kallay et al., 2006, Montcouquiol et al., 2006). Consequently, we performed pull-down assays using recombinant GST-fusions of Scribble PDZ1 to PDZ4 domains to examine binding of GFP-Vangl2 as well as β-PIX as a control. Whilst β-PIX bound to Scribble PDZ1 and PDZ3 as previously reported (Lim et al., 2017), we could also only detect binding of GFP-Vangl2 to PDZ2 and PDZ3 and this was dependent on the Vangl2 PBM (Figure 1).

**Table 1:** Summary of affinities of Vangl2, β-PIX and APC peptides for WT and mutant Scrib PDZ domains measured at pH 7.5 and 25 °C. NB denotes no binding, n.d. denotes not determined. Each of the value was calculated from at least three independent experiments. Values denoted by a and b were taken from (Lim et al., 2017) and (How et al., 2019), respectively.

|  | **K<sub>D</sub> (μM)** |  |  |  |
| --- | --- | --- | --- | --- |
| **SCRIB** | **Vangl2** | **mut Vangl2** | **β-PIX** | **APC** |
| PDZ1 WT | 24.49±3.90 | NB | 3.33±0.30<sup>a</sup> | 5.97±1.10<sup>b</sup> |
| PDZ1 H793A | 29.94±8.27 | n.d. | 62.77±22.43 | 18.80±6.16<sup>b</sup> |
| PDZ1 Q808H | 10.40±0.27 | n.d. | 3.11±0.1 | 6.06±0.65 |
| PDZ1 E814G | NB | n.d. | NB | NB |
| PDZ2 WT | 45.13±4.92 | NB | 67.84±7.90<sup>a</sup> | 35.94±1.10<sup>b</sup> |
| PDZ2 R896A | 26.74±4.01 | n.d. | 189.24±24.89 | 30.59±8.1 |
| PDZ2 H928A | NB | n.d. | NB | NB |
| PDZ3 WT | 40.16±3.92 | NB | 14.47±2.09<sup>a</sup> | 18.28±3.33<sup>b</sup> |
| PDZ3 P1043L | NB | n.d. | NB | NB |
| PDZ3 R1044Q | 20.93±4.67 | n.d. | 14.08±3.43 | 27.43±3.62 |
| PDZ4 WT | NB | NB | NB | NB |

**FIGURE 1:**
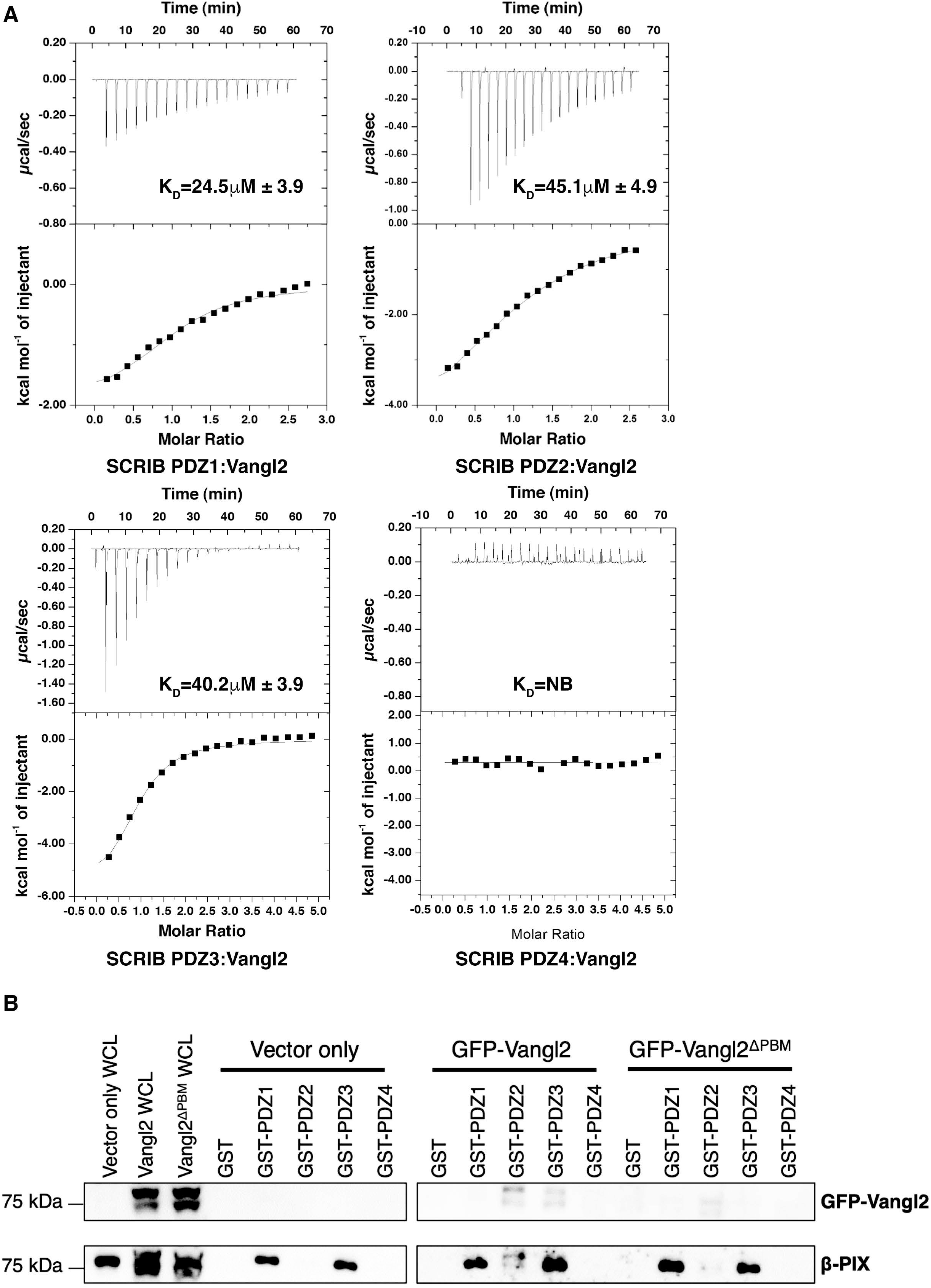
Interactions of Scribble PDZ domains with Vangl2. **A).** Binding profiles of isolated Scribble PDZ domains interaction with Vangl2 peptides are displayed. Each profile is represented by a raw thermogram (top panel) and a binding isotherm fitted with a one-site binding model (bottom panels). K_D_: dissociation constant; ±: standard deviation; NB: no binding. Each of the value was calculated from at least three independent experiments. **B)** GST-tagged individual Scribble PDZ domains 1-4 incubated with MCF10A lysates stably expressing vector only, GFP-Vangl2 and GFP-Vangl2^ΔPBM^. Bound proteins were recovered with glutathione resin and revealed by western transfer using anti-GFP and anti-β-PIX antibodies.

### Crystal structures of PDZ2:Vangl2 and PDZ3-Vangl2 complexes

To understand the structural basis and the mode of interaction for Scribble PDZ domains with Vangl2, we determined crystal structures of Scribble PDZ1:Vangl2, PDZ2-Vangl2 and PDZ3-Vangl2 as well as Scribble PDZ2 on its own (Figure 2, Table EV1, Figure EV2). As expected, all Scribble PDZ domains adopt the typical PDZ domain fold comprising a six-stranded β-sheet with two α-helices. All three Scribble PDZ domains comprise a compact globular fold featuring six β-strands and two α-helices that adopt a β-sandwich structure. The overall mode of Vangl2 peptide engagement is conserved across all three PDZ domain complexes, with PDZ1, PDZ2 and PDZ3 engaging Vangl2 in a manner where the peptide is bound in an anti-parallel orientation compared to the second β-strand whilst being sandwiched by the α2 helix in the canonical binding pocket of the PDZ domains. A comparison of the PDZ1 and PDZ3 domain structures on their own with their Vangl2 complex counterparts revealed no significant movement of secondary structure elements upon Vangl2 binding, as previously observed in complexes of PDZ1 and PDZ3 with β-PIX (Lim et al., 2017) and PDZ1 complexes with APC (How et al., 2019) and MCC (Caria et al., 2019). Similarly, superimposition of the Scribble PDZ2 domain structure with PDZ2:Vangl2 yielded an r.m.s.d of 0.453 A over 73 Cα atoms. A detailed examination of the Scribble PDZ1:Vangl2 complex (Figure 2 A) reveals that Vangl2 forms a number of direct contacts with the PDZ1 domain. The C-terminal carboxyl group of Vangl2 makes contacts with the main chains of L738^PDZ1^, G739^PDZ1^ and I740^PDZ1^, whilst the V521^Vangl2^ side chain is accommodated in a hydrophobic pocket formed by L738 ^PDZ1^, I740 ^PDZ1^, I742 ^PDZ1^, V797 ^PDZ1^ and L800 ^PDZ1^. Furthermore, hydrogen bonds between T519^Vangl2^:I742^PDZ1^, T519^Vangl2^:H793^PDZ1^, Q516^Vangl2^:T749^PDZ1^, E518^Vangl2^:S761^PDZ1^, E518^Vangl2^:R762^PDZ1^ are observed. In the Scribble PDZ2:Vangl2 complex (Figure 2 B), key features of the PDZ1:Vangl2 complex are conserved, with V521^Vangl2^ being located in a pocket formed by L872^PDZ2^, F874^PDZ2^, I876^PDZ2^, V932^PDZ2^ and L935^PDZ2^, whilst the C-terminal carboxyl group forms hydrogen bonds with the main chain of L872^PDZ2^, G873^PDZ2^ and F874^PDZ2^. Additional interactions are formed by S520^Vangl2^:S875^PDZ2^, T519^Vangl2^:H928^PDZ2^, as well as a salt bridge between E518^Vangl2^:S895^PDZ2^. In the PDZ3:Vangl2 complex (Figure 2 C), V521^Vangl2^ is engaged by a hydrophobic pocket formed by L1014^PDZ3^, L1016^PDZ3^, I1018^PDZ3^, V1075^PDZ3^ and L1079^PDZ3^ with the V521^Vangl2^ carboxyl group forming hydrogen bonds with the main chains of L1014^PDZ3^, G1015^PDZ3^ and L1016^PDZ3^. Other contacts are formed by T519^Vangl2^:I1018^PDZ3^, S520^Vangl2^:S1017^PDZ3^, T519^Vangl2^:H1071^PDZ3^, and E518^Vangl2^:S1039^PDZ3^.

**FIGURE 2:**
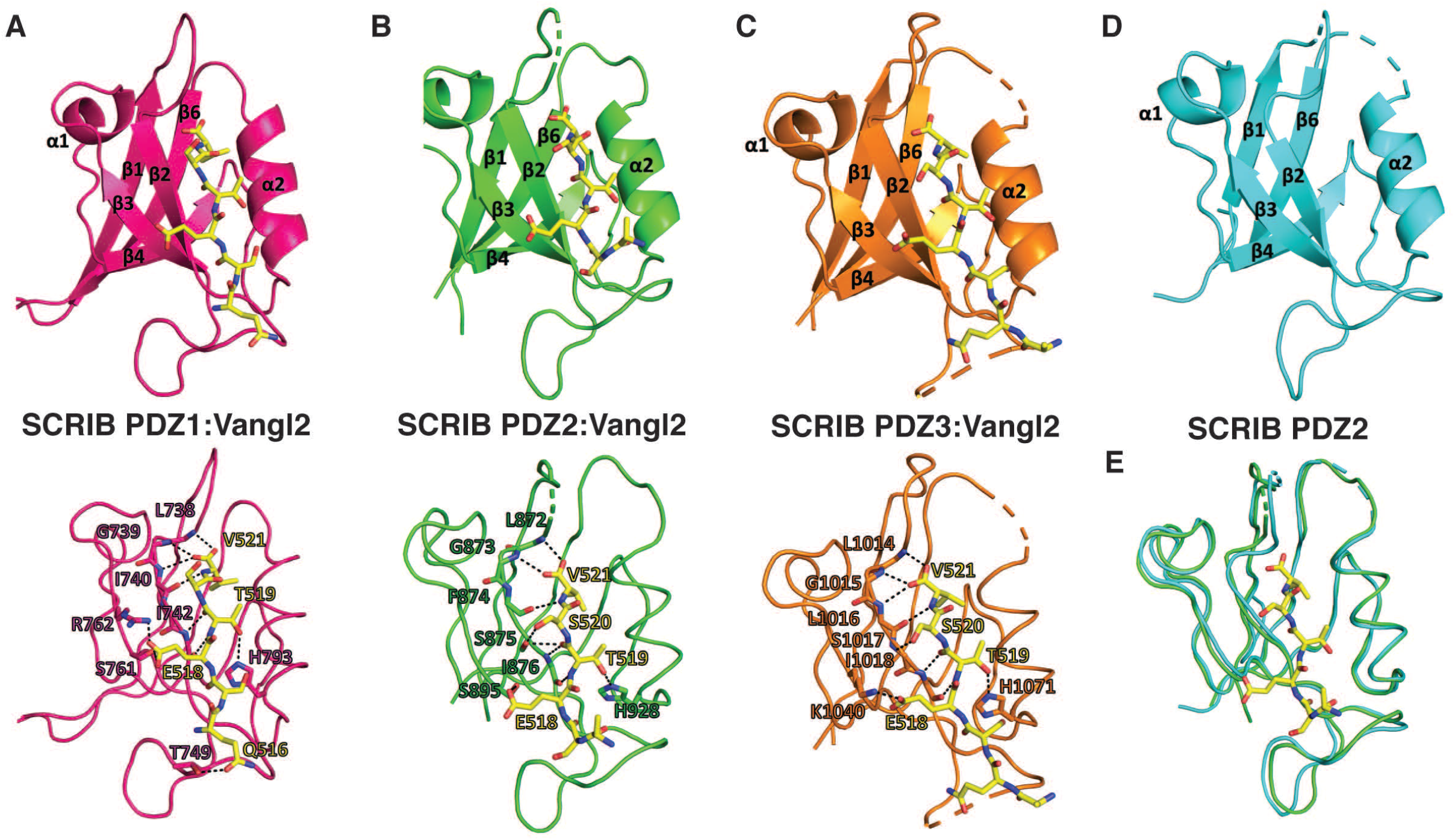
The crystal structures of PDZ1, PDZ2 and PDZ3 each bound to a Vangl2 peptide. The Vangl2 peptide engages individual PDZ domains via a shallow groove located between the β2 and α2. **A)** PDZ1 (magenta) is shown as a cartoon with Vangl2 peptide (yellow) represented as sticks. In the bottom panel, detailed interactions of Scribble PDZ1 with Vangl2 are shown as black dotted lines. Residues involved in these interactions are labelled. **B)** PDZ2 (green) is shown as a cartoon with Vangl2 peptide (yellow) represented as sticks. In the bottom panel, detailed interactions of Scribble PDZ2 with Vangl2 are shown as black dotted lines. Residues involved in these interactions are labelled. **C)** PDZ3 (orange) is shown as a cartoon with Vangl2 peptide (yellow) represented as sticks. In the bottom panel, detailed interactions of Scribble PDZ3 with Vangl2 are shown as black dotted lines. Residues involved in these interactions are labelled. **D)** Ligand-free PDZ2 is shown as a cartoon (cyan). **E)** Overlay of ribbon traces of PDZ2 with (green) and without Vangl2 peptide (forest green).

Whilst previous studies verified the mode of binding to Scribble PDZ1 and PDZ3 via mutagenesis (Lim et al., 2017, Caria et al., 2018), this has not been performed for Scribble PDZ2. Consequently, we generated a PDZ2 mutant where H928 was substituted by Ala (PDZ2H928A), which showed complete loss of binding to the Vangl2 PBM peptide (Figure 3). To validate our affinity measurements, we performed pull-down assays using recombinant GST-fusions of Scribble PDZ2 to GFP-Vangl2 and confirmed that PDZ2H928A could no longer bind to Vangl2 (Figure 3).

**FIGURE 3:**
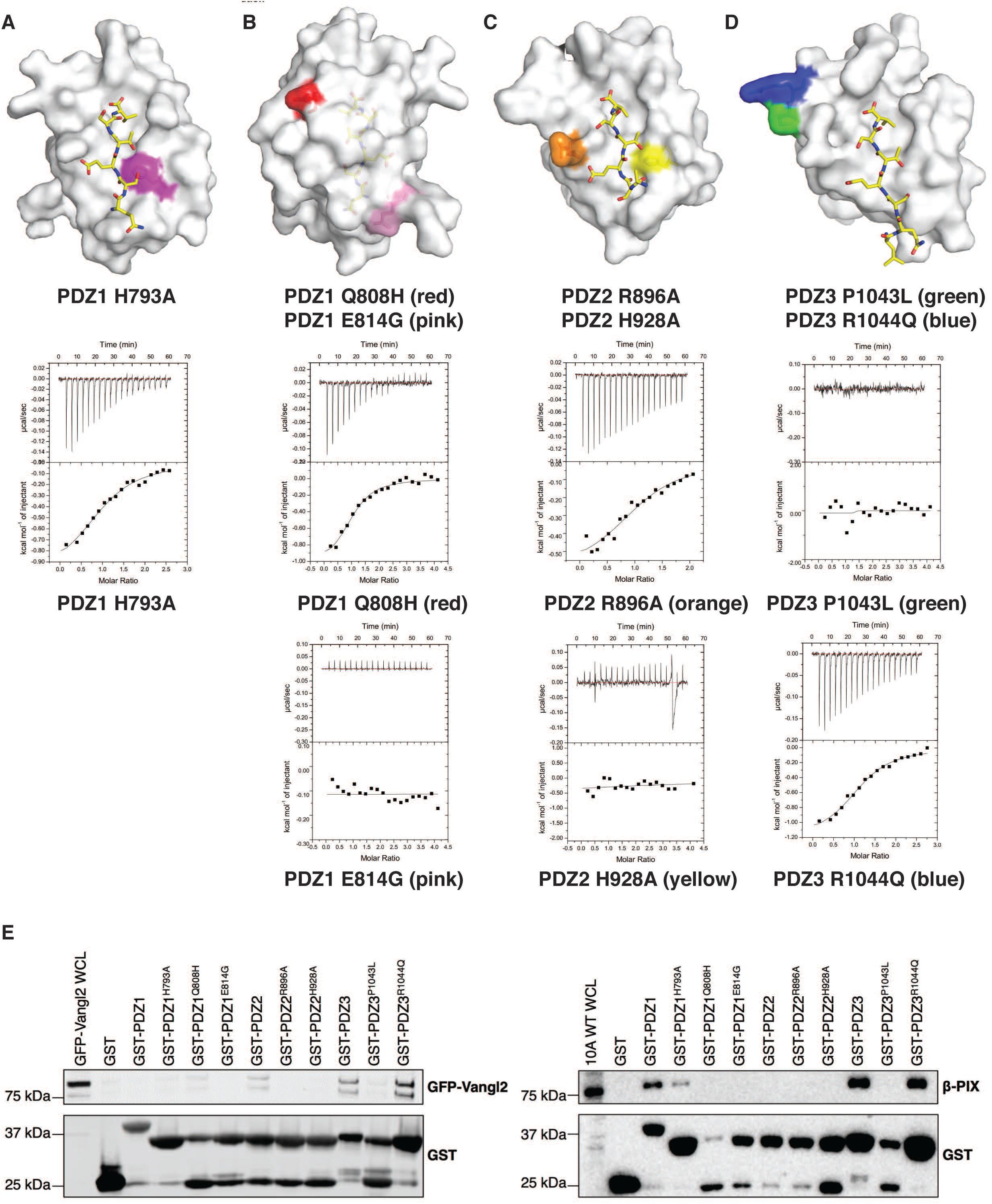
Interaction profiles of mutant Scribble PDZ with Vangl2 peptides. **A)** Location of Scribble PDZ domain mutations used in this study. Mutated residues are coloured on a gray surface representation of the relevant Scribble PDZ domain structure. Binding profiles of isolated mutant Scribble PDZ domain interactions with Vangl2 peptides. Each profile is represented by a raw thermogram (top panel) and a binding isotherm fitted with a one-site binding model (bottom panels). K_D_: dissociation constant; ±: standard deviation; NB: no binding. Each of the value was calculated from at least three independent experiments. **B)** GST-tagged Scribble PDZ domains, wildtype and with single-point mutations incubated with MCF10A WT lysates (left panel) and MCF10A cells stably expressing GFP-Vangl2 (right panel). Bound proteins were recovered with glutathione resin and revealed by western transfer using anti-GST, anti-β-PIX and anti-GFP antibodies.

In view of the reported physical and genetic interaction of Scribble and Vangl2 and their proposed role in coordinating neural tube closure in mice and humans, we then examined the ability of reported neural tube closure point-mutants in Scribble PDZ domains on their ability to bind the PBM of Vangl2 using ITC (Figure 3). The mutations examined identified in human neural tube closure defects were PDZ1Q808H (Kharfallah et al., 2017), PDZ3P1043L (Lei et al., 2013), and PDZ3R1044Q (Wang et al., 2018). We also engineered the equivalent mutation to the mouse *scrib1* ENU mutant, *crn2* (Stottmann et al., 2011) into human PDZ1, as PDZ1E814G, and examined engineered mutants PDZ1H793A, PDZ2R896A and PDZ2H928A. Scribble mutants PDZ1E814G and PDZ3P1043L revealed a loss of binding to Vangl2, whereas PDZ1Q808H maintained an affinity comparable to the wild-type interaction. Furthermore, despite its close proximity to PDZ3P1043L, PDZ3R1044Q maintained an affinity comparable to the wild-type interaction indicating the specificity of the PDZ3P1043L mutation on ligand binding. To confirm that the loss of binding of certain Scribble mutants was not due to an unfolding defect, we confirmed that all Scribble mutants were folded using circular dichroism spectroscopy (Figure EV3). To determine whether or not the mutations in Scribble PDZ domain mutants were specific for Vangl2 or acted more broadly we examined the ability of PDZ1E814G, PDZ1Q808H, PDZ3P1043L and PDZ3R1044Q as well as our engineered mutants for other well-known Scribble interactors, such as β-Pix (Lim et al., 2017) and APC (How et al., 2019). ITC measurements revealed that PDZ1E814G and PDZ3P1043L lost binding to both β-Pix and APC in addition to Vangl2. In contrast, PDZ1Q808H and PDZ3R1044Q bound interactors with affinities comparable to wild-type PDZ (Lim et al., 2017) and APC (How et al., 2019), with PDZ1Q808H binding β-Pix with 10.4 μM, APC with 6.0 μM and Vangl2 with 10.4 μM, whereas PDZ3R1044Q bound β-Pix with 14.1 μM, APC with 27.4 μM and Vangl2 with 20.9 μM. PDZ1H793A showed only a significant decrease in binding to β-Pix (62.77 μM), whereas PDZ2R896A showed decreased affinity to β-Pix and APC compared to wild-type Scribble PDZ2 binding (189.24 μM and 30.59 μM respectively). PDZ2H928A lost binding to all three interactors. To validate our affinity measurements we performed pull-down assays using recombinant GST-fusions of Scribble PDZ1 to PDZ4 domains to examine binding of GFP-Vangl2 as well as β-PIX as a control (Figure 3 E). We observed that PDZ3R1044Q, which displayed largely unchanged affinity to β-Pix, APC and Vangl2, was able to pull down β-Pix and Vangl2, similar to PDZ1Q808H. Other mutants including PDZ1E814G and PDZ3P1043L did not pull down either β-Pix or Vangl2. However, not all pull-down assays agreed with the measured affinities from ITC, with PDZ2R896A maintaining Vangl2 binding based on ITC but not in the pull-down assay.

## Discussion

Scribble is a large multi-domain scaffold protein that has been shown to play a pivotal role in cell polarity by integrating signaling from a multitude of interactors to control process such as cell migration and wound healing. This is predominantly achieved via Scribble’s four PDZ domains, which mediate the vast majority of Scribble interactions (Stephens et al., 2018). Whilst apicobasal cell polarity control is the most prominent of Scribble’s function, Scribble also plays an important role in the establishment and control of planar cell polarity (Milgrom-Hoffman and Humbert, 2018). The importance of Scribble function on planar cell polarity is illustrated by the impact of specific point mutations in Scribble, which manifest themselves as major developmental defects such as neural tube closure defects and disruption in the cochleae inner ear hair bundle orientation (Murdoch et al., 2003, Montcouquiol et al., 2003). Intriguingly, similar defects are observed for mutations in the transmembrane protein Vangl2, leading to the establishment of a genetic link between both proteins (Murdoch et al., 2001, Murdoch et al., 2003). Although interactions between Scribble and Vangl2 have previously been shown to involve Scribble’s PDZ2 and 3 domains, the detailed molecular and structural basis for these interactions remains to be clarified.

We now show that in addition to the previously identified interactions of Scribble PDZ2 and 3 domains with the C-terminal PBM of Vangl2, Scribble’s PDZ1 domain is also able to bind the Vangl2 PBM. Notably, affinities for all three interactions are comparable with a range of 24-45 μM K_D_, indicating that no distinct hierarchy in affinities exists for the Scribble:Vangl2 interactions. This is unusual in comparison with other Scribble interactions with e.g. β-PIX (Lim et al., 2017) or Gukh (Caria et al., 2018). Whilst Scribble also binds β-PIX with its PDZ1, 2 and 3 domains, PDZ1 is the highest affinity interactor with a K_D_ of 3.3 μM whereas PDZ2 is the lowest affinity site with a K_D_ of 67.8 μM (Lim et al., 2017). In contrast, Scribble engages tightly Gukh with its PDZ1 domain (K_D_=0.66 μM) whilst PDZ3 bound Gukh with only 27.8 μM, with no binding detected with its PDZ2 and 4 domains (Caria et al., 2018).

To understand the structural basis of Scribble’s PDZ domain interactions with the Vangl2 PBM, we determined complexes of Scribble PDZ1,2 and 3 domains with Vangl2. A comparison of the three PDZ domain complexes with Vangl2 reveals that whilst key family defining interactions are maintained and are near identical in all three including the engagement of the C-terminal Val (0) as well as the Thr in the −2 position, other interactions vary and offer scope for distinguishing between the domains. A comparison of PDZ1:Vangl2 with other determined Scribble PDZ1 complexes indicates that specific patterns can be seen depending on the nature of the specific PBM that is bound. For example, residues at the −5 position are shown to interact with the β2-β3 loop of PDZ1, whereas such interactions could not be observed in other Scribble PDZ domains. This behavior is also observed in *D. melanogaster.* This may be linked to the higher affinity of PDZ1 for its interactors compared to the other domains. A notable difference between PDZ1 and PDZ2 and 3 is that the latter utilized non-aromatic residues to engage the −5 position of bound PBM sequences, which preclude π-stacking as observed in Scribble PDZ1-βPIX (Lim et al., 2017) and *D.melanogaster* Scribble PDZ1-GukH (Caria et al., 2018). This supports the notion that the β2-β3 loop is a significant modulator of Scribble PDZ1 domain binding affinities, which is supported by mutagenesis data where β2-β3 loop mutations reduce binding affinity (Lim et al., 2017).

With the availability of high-resolution structures of all key Scribble interacting domains with Vangl2, we revisited the previously identified mutations in Scribble that lead to neural tube closure defects. Mapping of these point mutants (Milgrom-Hoffman and Humbert, 2018) indicates that in addition to mutations resulting in premature termination of Scribble, several mutations mapped to individual PDZ domains. Our analysis of these mutations using ITC revealed that although PDZ1Q808H, PDZ1E814G, PDZ3P1043L and PDZ3R1044Q are located distal from the canonical PDZ domain binding site, both PDZ1E814G and PDZ3P1043L are unable to bind to Vangl2 or indeed any other interactor tested including β-PIX and APC even though both mutants are folded, whereas PDZ1Q808H and PDZ3R1044Q displayed affinity that was comparable to the wild-type interaction. In contrast, a control mutant that mutated a key H928 residue in PDZ2 to Ala showed no binding. We note that a second engineered mutant, PDZ2R896A maintained binding to the Vangl2 PBM peptide when examined by ITC, but lost binding in pull-down assays with full-length Vangl2 as well as to β-PIX. Furthermore, whilst we were able to detect binding of the Vangl2 PBM peptide to Scrib PDZ1 and determine a crystal structure of the resultant complex, GST pull-down assays did not clearly detect this interaction as noted by others. This suggests that multiple factors may contribute to the interactions of Scribble PDZ domains with their interactors in addition to outright affinity, which impacts the ability to detect such interactions with a given approach.

The loss of binding of Vangl2 of two of the NTD associated mutants is intriguing when considering that both mutants were shown to be folded. We speculate that these results are due to secondary shell or other distal changes that indirectly impact the canonical ligand binding groove of PDZ domains, or affect PDZ domain stability in a manner that is not readily detectable by secondary structure changes and circular dichroism spectroscopy. An example for such a behavior was reported for the Tiam1 PDZ domain, which undergoes significant conformational changes on its secondary structures when interacting with different binding partners (Liu et al., 2013). Importantly, NMR and molecular dynamics approaches suggested that the β1- β2 and β3-α1 regions are dynamically linked, thus providing for a mechanism where changes distal to the canonical ligand binding site of the TIAM1 PDZ domain modulate ligand binding and interactions. In addition, allosteric regulation on the surface (Reynolds et al., 2011) of Scribble PDZ domains may be affected by the mutations we examined. PDZ domains such as PDZ3 from PSD97 have been shown to harbor sparse networks of coevolving amino acids, which when mutated may substantially impact ligand binding behaviour (Chi et al., 2008, McLaughlin et al., 2012). Whilst comparable analyses to PSD97 PDZ3 have not been performed for Scribble PDZ domains, considering that the networks of coevolving amino acids in PSD97 are tightly linked to the structure and function of the PDZ domain it seems plausible that analogous networks may also be found in Scribble PDZ domains.

Interestingly, an analysis of our Scribble PDZ domain structures using mCSM (Pires et al., 2014) revealed that PDZ1E814G, PDZ3P1043L and PDZ3R1044Q were all predicted to be destabilizing mutations that would result in a reduced affinity for an interactor, whereas PDZ1Q808H was predicted to be a stabilizing mutation with increased affinity for an interactor. Nevertheless, the observed loss of binding of Scribble PDZ1E814G, PDZ3P1043L mutants allows the rationalization of the NTD phenotype, which would be due to a loss of binding to Vangl2. However, considering the loss of binding of PDZ1E814G and PDZ3P1043L for APC and β-PIX in addition to Vangl2 suggests that these mutations are not specific for Vangl2, and that the neural tube defect phenotype arises due to the complete loss of interaction functionality of the affected Scribble PDZ domain. This would be consistent with the neural tube closure defect phenotype of mice with truncating mutations of Scribble with complete loss PDZ3 and PDZ4 (*crc*, 947fs mutation, (Murdoch et al., 2003)) although this mutation also destabilizes the Scribble protein itself (Moreau et al., 2010, Yates et al., 2013).

In summary, we showed that biochemically Scribble is able to bind the Vangl2 PBM using its PDZ1,2 and 3 domains. Furthermore, we determined the structural basis for Vangl2 binding for all the interacting Scribble PDZ domains, and establish that well known NTD mutants of Scribble outside the canonical ligand binding groove affect its PDZ domain ligand binding behaviour, with mutant PDZ domains unable to engage a broad set of known Scribble interactors and not only Vangl2. Our findings raise the possibility that Scribble mutations associated with diseases such as NTDs are not disease causing due to loss of a single interaction, and instead support a view where a given mutation perturbs the overall interaction network of the adaptor protein Scribble by impacting multiple interactions and the dynamic interplay of different interactor with Scribble. Overall these findings form a mechanistic platform to understand how dysregulation of the Scribble interactions impacts planar cell polarity establishment, and may cause tissue disruption that manifests itself as neural tube closure defects.

## Materials and methods

### Protein expression and purification

Recombinant human Scrib (Uniprot accession number: Q14160) domains spanning PDZ1 (728–815); PDZ2 (860–950); PDZ3 (1002–1094); PDZ4 (1099–1203)) as well as Scribble mutants were expressed using *Escherichia coli* BL21 (DE3) pLysS cells (BIOLINE) as Glutathione S-transferase or Maltose Binding Protein fusions (Rautureau et al., 2012) and purified as previously described (Lim et al., 2017). Scribble PDZ domain mutants PDZ1 H793A, PDZ1 Q808H, PDZ1 E814G, PDZ2 R896A, PDZ2 H928A, PDZ3 P1043L and PDZ3 R1044Q were synthesized as codon-optimized synthetic dDNA and cloned in the pGEX-6P3 vector (Genscript). Mutant Scribble PDZ domains were expressed and purified as previously described (Lim et al., 2017).

### Isothermal titration calorimetry

Purified wild type and mutant human Scrib PDZ domains were used in isothermal titration experiments against 8-mer peptides spanning the C terminus of human Vangl2 (Uniprot accession number: Q9ULK5; RLQSETSV) or a non-binding mutant of Vangl2 (RLQSEASA) to determine the affinity for Scribble PDZ domains. Titrations were performed at 25°C with a stirring speed of 750 rpm using the MicroCalTM iTC200 System (GE Healthcare). A total of 20 injections with 2 μL each and a spacing of 180 seconds were titrated into the 200 μL protein sample (25 mM Hepes pH 7.5, 150 mM NaCl), except for the first injection which was only 0.4 μL. Protein concentration of 75 μM against peptide concentration of 0.9 mM were used. Peptides were purchased from Genscript (San Francisco, CA, USA). Raw thermograms were processed with MicroCal Origin^®^ version 7.0 software (OriginLabTM Corporation) to obtain the binding parameters of each interaction. A synthetic pan-PDZ binding peptide referred to as superpeptide (RSWFETWV) was used as a positive control (Caria et al., 2018).

### Protein crystallisation, data collection and refinement

PDZ1, 2 and 3 domain proteins were mixed with wild-type Vangl2 peptide at a 1:20 molar excess to reconstitute the relevant complexes. Samples were subjected to crystallization screening in-house using a Gryphon nanodispenser (Art Robbins Instruments). PDZ1:Vangl2 crystals were obtained at 25 mg/mL in 0.1M Phosphate/citrate pH 4.2 and 40% (v/v) PEG 300. Both PDZ2 and PDZ2-Vangl2 complex crystals were obtained at 15mg/ml in 0.2M Potassium thiocyanate and 20% (w/v) PEG 3350. PDZ3-Vangl2 crystals were obtained at 10mg/ml in 0.1 M HEPES pH 7.8 and 66% (v/v) MPD.

### Diffraction data collection and structure determination

All crystals were flash cooled in mother liquor, the former supplemented with 20% (v/v) ethylene glycol (PDZ2-Vangl2 and PDZ1-Vangl2 crystals). All diffraction data were collected on the MX1 beamline at the Australian Synchrotron equipped with an ADSC Quantum 210r CCD detector (Area Detector Systems Corporation, Poway, California, USA) or the MX2 beamline equipped with the EIGER 16M detector. Diffraction data were integrated using *XDS* (Kabsch, 2010), followed by *AIMLESS* (Winn et al., 2011) for merging and scaling. The structure of the PDZ1-Vangl2 complex was solved by molecular replacement using *Phaser* (McCoy, 2007) using the structure of Scribble PDZ1 (PDB code 6MTV (Caria et al., 2019)) as a search model. The structure of the PDZ3-Vangl2 complex was solved by molecular replacement using *Phaser* (McCoy, 2007) using the structure of Scribble PDZ3 (PDB: 4WYT) as a search model. The structure of the PDZ2 domain on its own was also solved using the final model of the PDZ3-Vangl2 structure. The PDZ2-Vangl2 complex was then subsequently solved using the final model of the PDZ2 domain structure. The solutions produced by *Phaser* were manually rebuilt over multiple cycles using *Coot (Emsley and Cowtan, 2004)* and refined using *PHENIX* (Afonine et al., 2012). All images were generated using the PyMOL Molecular Graphics System, Version 1.8 Schrödinger, LLC. All software was accessed using the SBGrid suite (Morin et al., 2013).

### Circular dichroism spectroscopy

Proteins were buffer exchanged into 15 mM Na_2_HPO_4_/NaH_2_PO_4_ buffer. Spectra were acquired a protein concentrations ranging from 0.080 - 0.150 mg/ml on an AVIV 420 CD spectrometer at 25℃. Wavelength scans were performed from 190 to 260 nm at 1 nm intervals and an averaging time of 4 seconds. Data were processed using the AVIV Biomedical software and displayed using Excel.

### Cell culture

MCF10A cells were cultured in Dulbecco’s Modified Eagle Medium:F12 (DMEM:F12) supplemented with 5% donor horse serum (Gibco), 10 μg/mL insulin (Actrapid, Novo Nordisk Pharmaceuticals Ltd.), 0.5 μg/mL hydrocortisone (Merck), 20 ng/mL epidermal growth factor (Peprotech), 100 ng/mL cholera toxin (List Biological Laboratories Inc.), 100 units/mL penicillin, and 100 μg/mL streptomycin and maintained at 37°C in 5% CO_2_.

### Plasmid construct and viral transduction

Wild-type Vangl2 cDNA and Vangl2 cDNA missing the last four amino acids (ETSV) were cloned into MSCV vector using Gibson Assembly to generate N-terminally fused GFP-Vangl2 and GFP-Vangl2^ΔPBM^ expression constructs. Retroviral particles were produces by co-transfection of Hek293T cells with retroviral packaging plasmids and MSCV-based retroviral vectors expressing GFP-Vangl2, GFP-Vangl2^ΔPBM^ and the MSCV vector only and subsequently infecting low-passage MCF10A cells for stable expression. Infected cells were selected with 2 μg/mL puromycin for 1 week.

### GST pulldowns

GST-fusions of wild type and mutant Scribble PDZ domains were expressed and purified as described above. MCF10A cells were lysed in NETN lysis buffer (20 mM Tris-Cl, pH8.0; 100 mM NaCl; 1 mM EDTA; 0.5% Nonidet P-40) supplemented with a phosphatase inhibitor and protease inhibitor cocktail (Roche Diagnostics) for 10 minutes. 250 μg of MCF10A cell lysate and 5 μg of individual GST-tagged recombinant proteins were mixed and incubated with glutathione Sepharose 4b beads (GE Healthcare) at 4°C overnight. Beads were washed three times with NETN buffer prior to being eluted in loading buffer.

### Western Blot

Eluted samples were resolved on standard sodium dodecyl sulphate-polyacrylamide gels and transferred to polyvinylidene difluoride membrane (Millipore). Membranes were blocked in 3% BSA in 0.2% Tween-20 in PBS for 1 hour, then incubated with appropriate primary and secondary antibodies in blocking solution. Membranes were visualised using an Odyssey infrared imager (LiCor Biosciences). Antibodies used were rabbit polyclonal anti-GFP antibody (A-6455; Invitrogen); rabbit polyclonal anti-β-PIX antibody (4515S; Cell Signalling Technology) and rabbit polyclonal anti-GST antibody (71-7500; Invitrogen).

## ACKNOWLEDGEMENTS

We thank staff at the MX beamlines at the Australian Synchrotron for help with X-ray data collection, and the CSIRO C3 Collaborative Crystallization Centre for assistance with crystallisation and the Comprehensive Proteomics Platform at La Trobe University for core instrument support. This research was undertaken in part using the MX2 beamline at the Australian Synchrotron, part of ANSTO, and made use of the ACRF detector. This work was supported in whole or part by the National Health and Medical Research Council Australia (Project Grant APP1103871 to MK, POH; Senior Research Fellowship APP1079133 to POH), Australian Research Council (Fellowship FT130101349 to MK) and La Trobe University (Research Focus Area Understanding Disease grant 2000002510).

## AUTHOR CONTRIBUTIONS

JH: Acquisition of data; Analysis and interpretation of data; Drafting and revising the article. RKS: Acquisition of data; Analysis and interpretation of data; Drafting and revising the article.

KYBL: Acquisition of data; Analysis and interpretation of data.

POH: Conception and design; Analysis and interpretation of data; Drafting and revising the article

MK: Conception and design; Acquisition of data; Analysis and interpretation of data; Drafting and revising the article

## Data availability statement

Coordinate files have been deposited in the Protein Data Bank under the accession code 6XA6, 6XA7, 6XA8 and 7JO7 . Raw diffraction images were deposited on the SBGrid Data Bank (Meyer et al., 2016) using their PDB accession numbers.

## Expanded view figure legends

**Table EV1:**
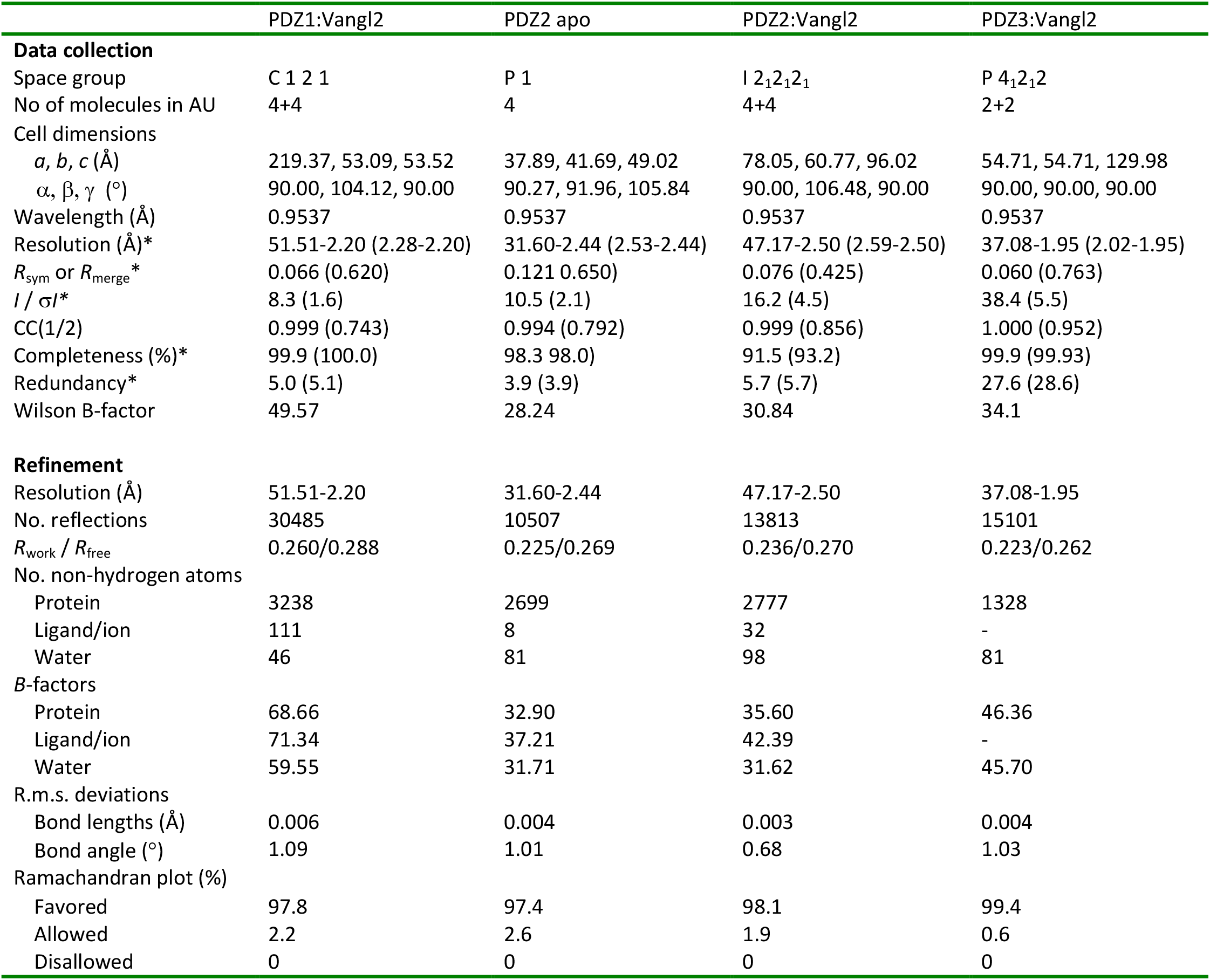
X-ray crystallographic data collection and refinement statistics.

**FIGURE EV2:**
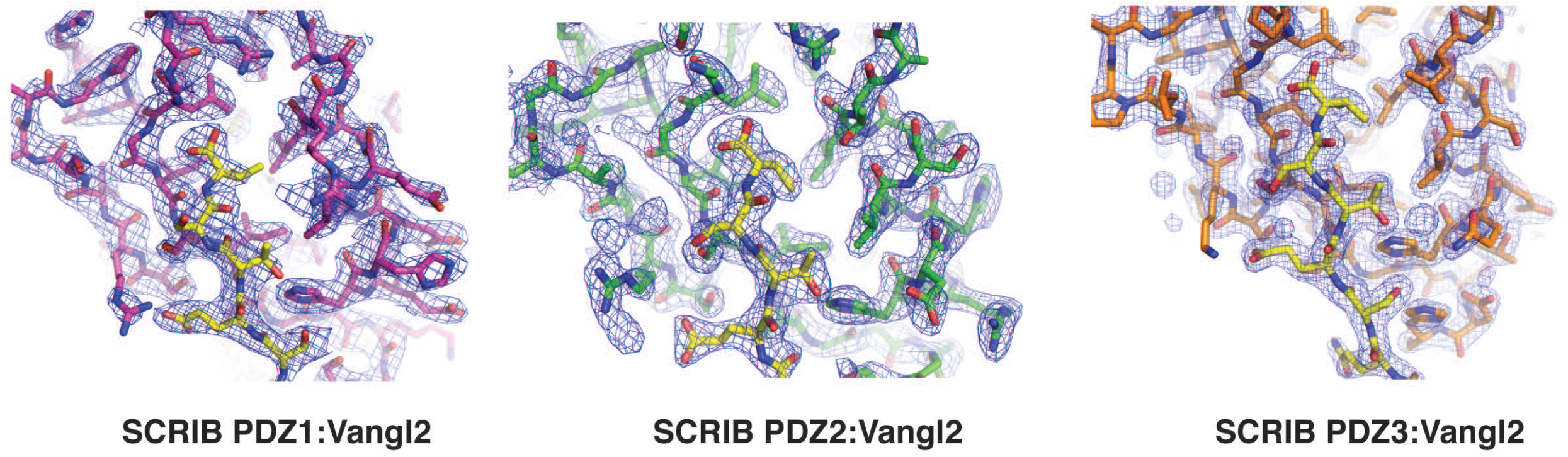
2Fo-Fc electron density maps of PDZ1, PDZ2 and PDZ3 complexes with Vangl2 as well as PDZ2 on its own. **A)** Electron density map encompassing the binding groove of Scribble PDZ1 (magenta sticks) in complex with Vangl2 peptide (yellow sticks). The electron density map is shown as a blue mesh contoured at 1.5 σ. **B)** Electron density map encompassing the binding groove of Scribble PDZ2 (green sticks) in complex with Vangl2 (yellow sticks). The electron density map is shown as a blue mesh contoured at 1.5 σ. **C)** Electron density map encompassing the binding groove of Scribble PDZ3 (orange sticks) in complex with Vangl2 peptide (yellow sticks). The electron density map is shown as a blue mesh contoured at 1.5 σ.

**FIGURE EV3:**
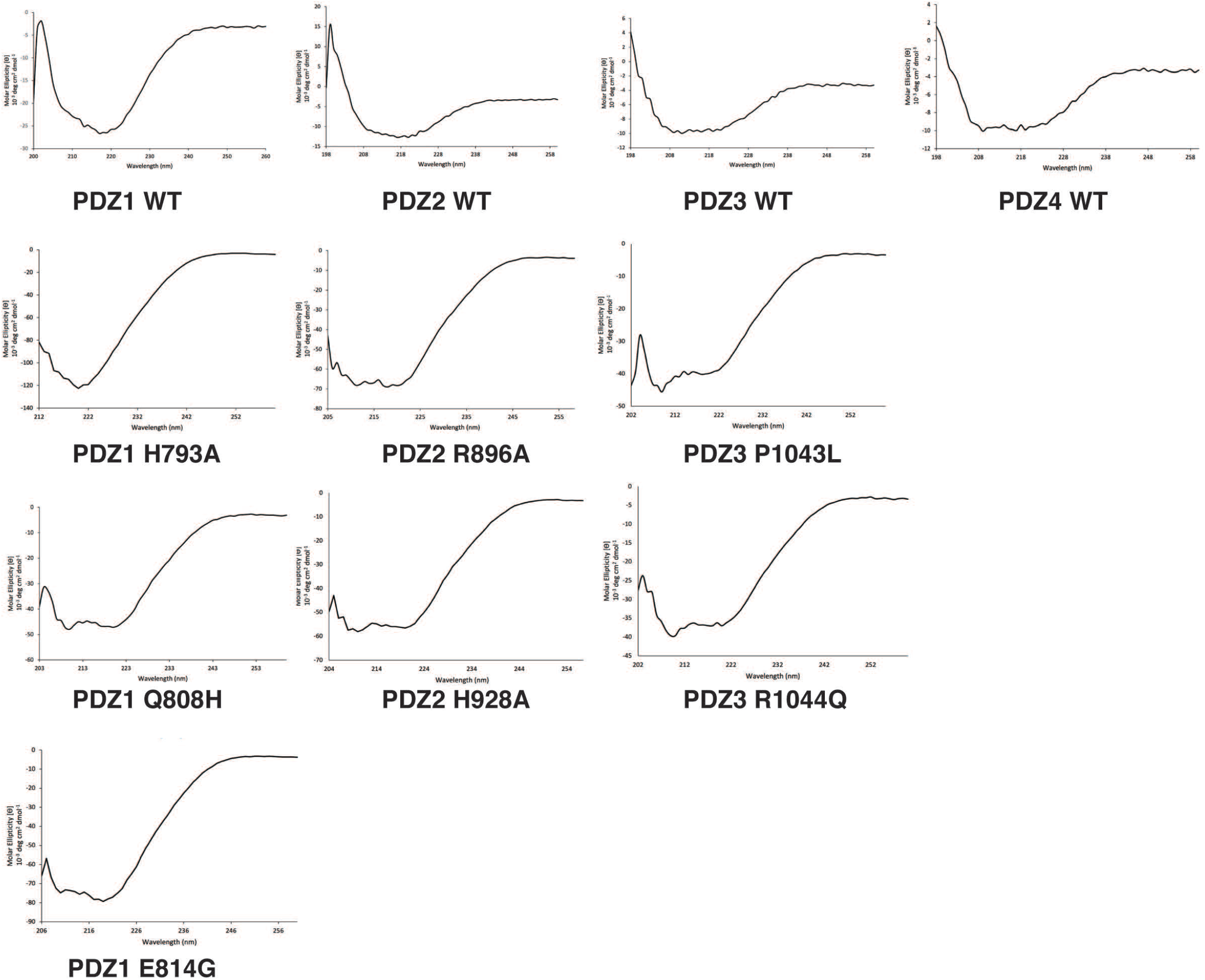
Circular dichroism spectroscopy of wild type and mutant Scribble PDZ1, PDZ2, PDZ3 and PDZ4 domains. Circular dichroism spectra recorded for wild type and mutant Scribble PDZ1,2,3 and 4 domains indicated that there were no major spectral differences between the proteins, suggesting that they were similarly folded, with mutations not leading to unfolding of the PDZ domains.

